# SET-M33 peptide as a selective *in vitro* antimicrobial agent against the porcine respiratory pathogen *Glaesserella parasuis*

**DOI:** 10.1101/2025.11.05.686722

**Authors:** Ana Laura Pereira Lourenço, Oussama Rouam el Khatab, Chiara Falciani, Alessandro Pini, Virginia Aragon, Marta Cerdà-Cuéllar, Karl Kochanowski

## Abstract

As we face the threat from bacterial pathogens that are resistant to many conventional antibiotics, many current research efforts focus on expanding our arsenal of antimicrobial compounds. However, identifying use cases in which such new antimicrobials can effectively target pathogens while minimizing collateral damage in the commensal microbiota remains a challenge. To tackle this challenge, we focused on one new antimicrobial, the synthetic antimicrobial peptide SET-M33, and examined its ability to target porcine respiratory pathogens and a collection of porcine commensal nasal microbiota members *in vitro*. Our experiments revealed three key results. First, there were large differences in SET-M33 sensitivity across the tested strains. In particular, pathogenic *Glaesserella parasuis* was highly sensitive to SET-M33 at concentrations that did not affect the growth of most commensal strains. Second, some of the tested commensal strains (i.e. *Rothia nasimurium* and *Staphylococcus aureus*) were able to inactivate SET-M33 during *in vitro* cultivation. Third, despite this potential for SET-M33 inactivation by commensal strains, SET-M33 was still able to selectively eliminate pathogenic *G. parasuis* from *in vitro* co-cultures that also contained *R. nasimurium*. Overall, this study highlights the substantial complexity that emerges from the interplay between antimicrobials, pathogens, and commensals, even within a comparatively simple *in vitro* system, and provides a template for identifying suitable use cases for newly developed antimicrobials.

## Introduction

Antimicrobial resistance (AMR) in pathogenic bacteria is among the most pressing global health burdens we currently face, and the number of AMR-related deaths have been predicted to rise to 10 million per year by 2050, with tremendous added economic impact (1–3). While advances in vaccination, sanitation, and disinfection provide new avenues for preventing the spread of pathogenic bacteria, such measures are unlikely to eradicate infectious diseases entirely. Therefore, having a diverse arsenal of effective antimicrobials that work against current (and future) bacterial pathogens will be fundamental. Of particular importance will be the development of antimicrobials with alternative modes of action (4–6), which can circumvent resistance mechanisms that are already prevalent in clinical isolates (7–9).

Towards the development of new antimicrobials with alternative modes of actions, antimicrobial peptides (AMPs) have emerged as particularly promising candidates (10,11). They are naturally occurring molecules that can be found as part of the immune response of all living organisms. Apart from the diversity of AMPs found in nature, they can also be made synthetically (4,12–15). Synthetic AMPs offer notable advantages over naturally isolated counterparts, particularly in addressing limitations related to peptide stability and toxicity. Advances in machine learning and AI-guided design are driving the development of computational platforms that enable the efficient discovery and optimization of AMPs with improved therapeutic properties (16–19). In general terms, AMPs present various mechanisms of action with non-specific targets (20), thus facilitating the effective killing of a broad range of pathogens including multi-drug resistant isolates (21–25). Moreover, AMPs have been shown to be less prone to acquired resistance (8,26,27), even though resistance to AMPs can occur (28,29). Thus, antimicrobial peptides represent a versatile class of therapeutics with highly tunable physicochemical and functional properties. However, their optimal application remains challenging, as clear strategies for aligning specific AMP characteristics with appropriate clinical use cases are still lacking.

In this study, we focus on one such synthetic AMP, namely SET-M33, which was originally selected in a random phage library against *Escherichia coli* and was later rationally modified to have a tetra-branched configuration (30). SET-M33 has already been shown to be effective against bacterial pathogens causing disease in humans, including *Klebsiella pneumoniae*, *Pseudomonas aeruginosa*, *Acinetobacter baumannii*, and *Staphylococcus aureus* (30–33). However, whether this AMP is effective also against other pathogens – and in particular non-human pathogens causing severe disease in production animals – remains to be clarified. Moreover, the potential impact of SET-M33 on the commensal microbiota (which constitutes the first line of defense against many respiratory and intestinal pathogens) has not yet been investigated.

Here, we tackled these questions using the porcine nasal microbiota as our model system and a three-pronged approach. First, we examined the effectiveness of SET-M33 against strains representing three of the most important bacterial pathogens causing respiratory disease in piglets, namely *Streptococcus suis, Actinobacillus pleuropneumoniae* and *Glaesserella parasuis*. *In vitro* testing against reference strains revealed large differences in susceptibility across bacterial species. In particular, we found that a virulent *G. parasuis* strain was highly sensitive to SET-M33 treatment. Second, we leveraged a recently developed collection of porcine nasal microbiota strains (34) to examine the impact of SET-M33 on the commensal microbiota. We found large differences in susceptibility against this AMP, with some commensal microbiota members being unaffected even by high SET-M33 doses. Third, we demonstrated that these differences in susceptibility between virulent *G. parasuis* and commensal microbiota members can be exploited to specifically eradicate the pathogen from pathogen-commensal co-cultures. Overall, this study highlights the potential of SET-M33 as an effective new therapeutic against *G. parasuis*, and more generally provides a template for the identification of use cases for newly developed antimicrobials.

## Results

### Section 1: Impact of SET-M33 on different porcine respiratory pathogens

As our case study of how a synthetic antimicrobial such as SET-M33 could be used to combat animal infections, we decided to focus on bacterial pathogens that infect the respiratory tract of piglets. This decision was motivated by the prevalence of such infections in commercial pig production, and by the increased pressure from regulators to develop alternatives to conventional antibiotics in animal production. Specifically, we focused on three pathogen species, namely *Streptococcus suis*, *Actinobacillus pleuropneumoniae* and *Glaesserella parasuis*, which account for a large portion of respiratory infections in piglets. For each species, we selected a highly virulent reference strain (i.e. *S. suis* P1/7, *A. pleuropneumoniae* 4074 and *G. parasuis* Nagasaki) and challenged it with varying concentrations (0.62 to 20 µM, matching the effective *in vivo* concentration range, (35,36)) of either SET-M33, or its D-stereoisomer SET-M33D (37).

We found large differences in susceptibility across the tested strains (**Figure 1**): *S. suis* P1/7 and *A. pleuropneumoniae* 4074 were largely insensitive to both SET-M33 and SET-M33D (minimal inhibitory concentration (MIC) of 20 µM or higher), whereas *G. parasuis* Nagasaki was highly sensitive to both compounds (MIC < 3 µM).

**Figure 1.**
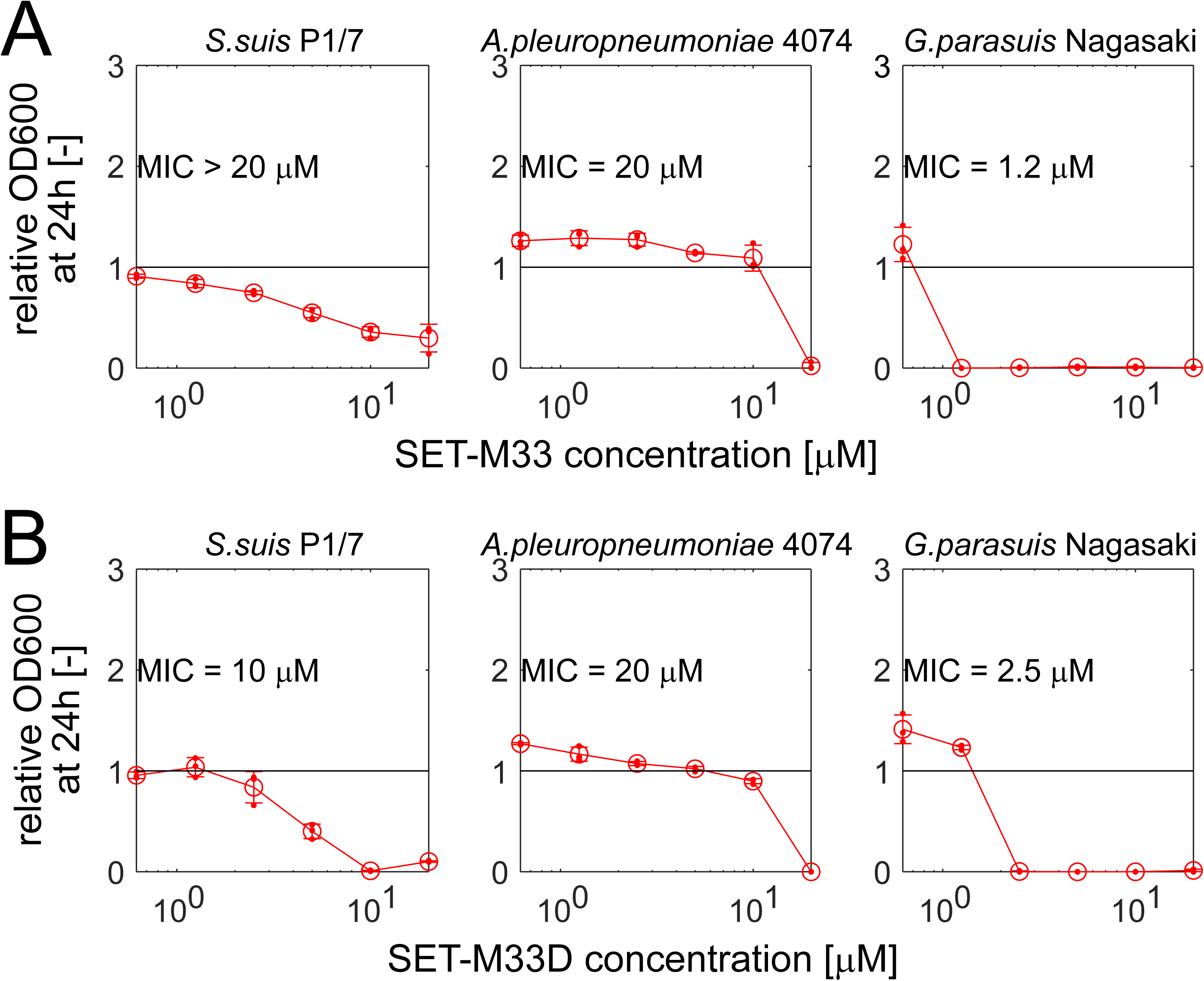
Impact of antimicrobial peptides SET-M33 (A) and SET-M33D (B) on three porcine respiratory pathogens. Relative OD600 (relative to untreated controls) after 24h incubation in planktonic BHI+ cultures with different compound concentrations for each strain. Error bars denote standard deviation (n = 3), small circles denote individual replicate wells. In each case, the MIC was determined analytically as the lowest compound concentration with a relative OD600 of <10%, and confirmed by visual inspection of the growth plates.

### Section 2: impact of SET-M33 on commensal nasal microbiota

In addition to effectively targeting pathogenic bacteria, an ideal antimicrobial peptide should also preserve the host’s commensal microbiota as much as possible, since such collateral damage has been associated with an increased susceptibility to subsequent infections (38–40). To test the potential of SET-M33 to cause such collateral damage, we made use of a recently developed porcine nasal consortium (PNC8), a collection of strains which represent the most *in vivo* abundant genera in the nasal microbiota of healthy piglets (34). For the six PNC8 strains that grew robustly in the tested condition (BHI+, the same condition as used for the pathogenic strains), we tested their susceptibility to both SET-M33 and SET-M33D in planktonic cultures using the same approach as described above.

We again observed large differences in susceptibility across the tested strains (**Figure 2**), which ranged from high sensitivity to both compounds (*Lactobacillus plantarum* KD9-5, *Neisseria shayeganii* GM3-3) to low sensitivity to either one (*Rothia nasimurium* UK1-9) or both (*Streptococcus pluranimalium* LG3-6, *Staphylococcus aureus* EJ41-2) compounds. Interestingly, in some cases (especially for *R. nasimurium* UK1-9) we observed an increase in OD600 compared to untreated controls at low AMP concentrations, which was robust across repeat experiments (**Supplementary Figure 1**). It is unlikely that this OD600 increase is caused by the AMP being catabolized for biomass production (since the tested compounds were being used at low micromolar concentrations), but we currently do not have any alternative explanations.

**Figure 2.**
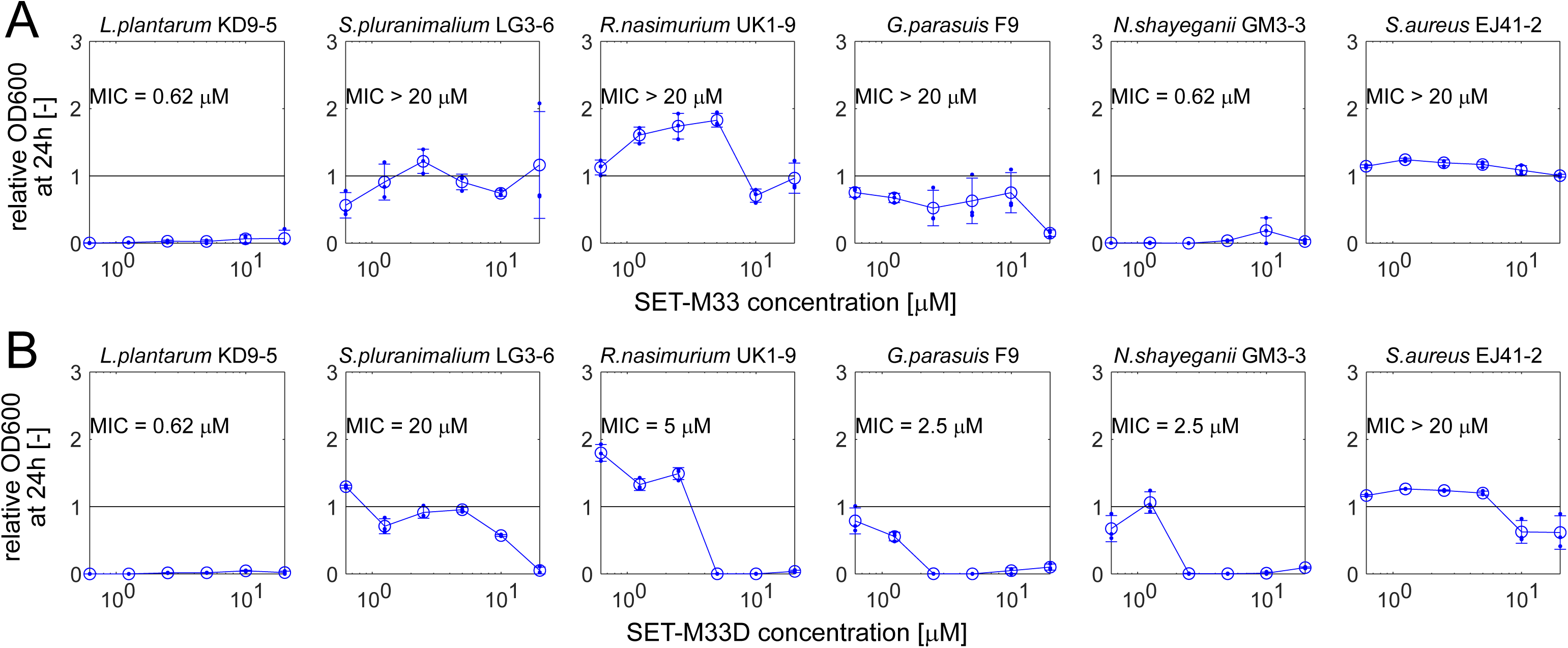
Impact of antimicrobial peptides SET-M33 (A) and SET-M33D (B) on commensal porcine nasal microbiota members. Relative OD600 (relative to untreated controls) after 24h incubation in planktonic BHI+ cultures with different compound concentrations for each strain. Error bars denote standard deviation (n = 3), small circles denote individual replicate wells. In each case, the MIC was determined analytically as the lowest compound concentration with a relative OD600 of <10%, and confirmed by visual inspection of the growth plates.

When comparing the MIC values of pathogenic and commensal strains for the two tested compounds (**Figure 3**), we found that in particular for SET-M33, there is a substantial difference in MIC between the pathogenic *G. parasuis* strain Nagasaki (MIC = 1.25 µM) and most tested PNC8 strains (MIC at or above 20 µM). Moreover, our data suggested that this difference in MIC could not simply be explained by a differential sensitivity to SET-M33 between a Gram-negative pathogen and Gram-positive commensals (the most sensitive commensal, *L. plantarum* KD9-5 is also Gram-positive).

**Figure 3.**
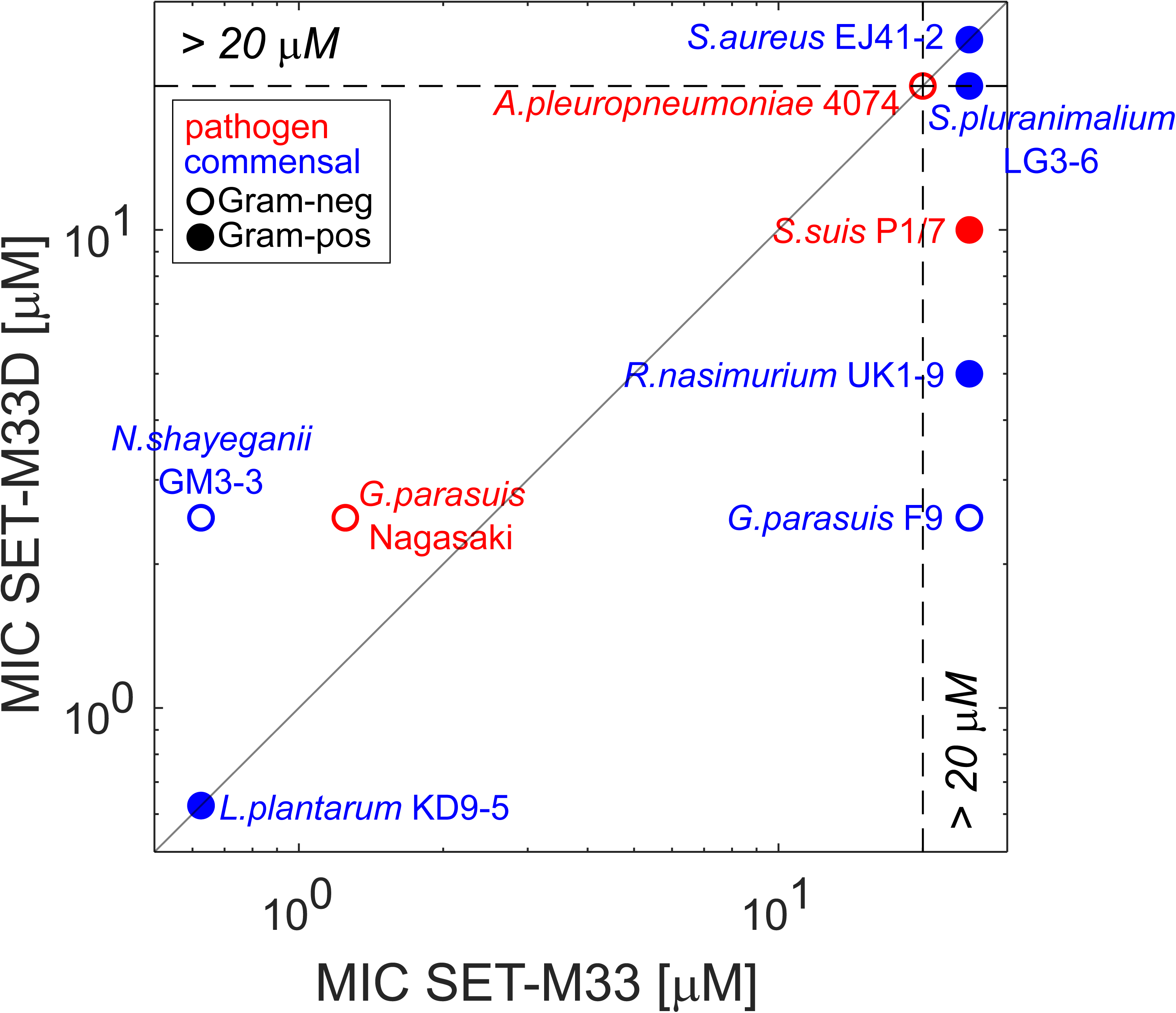
Comparison of MICs for SET-M33 (x-axis) and SET-M33D (y-axis). MICs were determined as described before (i.e. in each case, the MIC was determined as the lowest compound concentration with a OD600 relative to untreated controls of <10%). Red: pathogenic strains. Blue: commensal strains. Full circles denote Gram-positive strains, empty circles denote Gram-negative ones. See **Supplementary** Figure 1 for day-to-day reproducibility of MIC quantification.

### Section 3: Examining potential SET-M33 degradation by drug-tolerant commensal strains

The experiments described above revealed large differences in susceptibility to SET-M33 not only among pathogenic strains (**Figure 1**), but also among commensal nasal microbiota members (**Figure 2**). In particular, the data suggested that a large fraction of tested commensal strains can tolerate high doses of at least one of the tested SET-M33 variants (see Figure 3), which was unexpected given the previously described broad activity range of these compounds against many diverse bacterial species (31,33,37,41).

Hence, we wanted to examine the underlying mechanism of SET-M33 tolerance in the commensal strains. Given that SET-M33 (like many antimicrobial peptides) has previously been shown to be susceptible to degradation by cellular proteases (37), we reasoned that the most parsimonious tolerance mechanism would be the inactivation of the compound by these commensal strains. To test this conjecture, we designed a two-step cultivation experiment, in which SET-M33 is first incubated overnight with cultures containing highly tolerant commensal strains, and then its residual activity in the spent media is tested against a sensitive “sentinel” strain (see schematic in **Figure 4A**).

**Figure 4.**
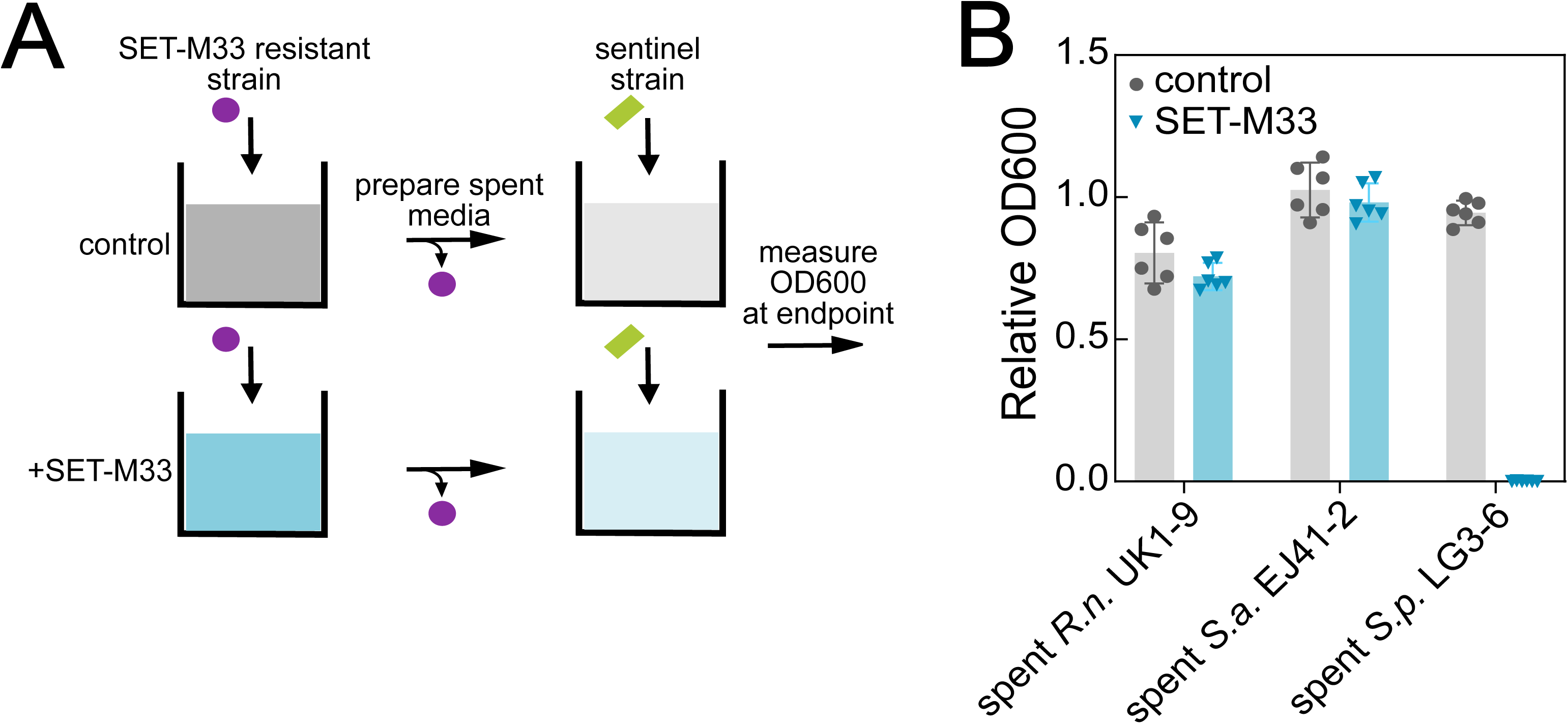
Evaluating the residual activity of SET-M33 after incubation with tolerant commensal nasal microbiota members. **A)** Schematic of approach. **B)** Relative OD600 (compared to fresh BHI+ cultivation media) of a SET-M33-susceptible sentinel strain (*Escherichia coli* NCM3722) cultivated in spent media obtained from commensal nasal microbiota members incubated with 10 microM SET-M33 overnight, and then mixed 1:1 with fresh media (to replenish depleted growth nutrients). Error bars denote standard deviation (n = 6, from two independent experiments), circle denote individual replicate cultures. *R.n*., *Rothia nasimurium*; *S.a*., *Staphylococcus aureus*; S.p., *Streptococcus pluranimalium*.

We found that in two of the commensal strains tested, namely *R. nasimurium* UK1-9 and *S. aureus* EJ41-2, the SET-M33 sensitive sentinel strain was able to grow in their respective spent media, suggesting that SET-M33 activity was indeed lost (**Figure 4B**, blue bars). In contrast, in the third tested commensal, *S. pluranimalium* LG3-6, SET-M33 retained its activity in the spent media. Control experiments showed that prolonged incubation of SET-M33 in bacteria-free BHI+ media did not affect its activity (**Supplementary Figure 2**). Thus, these data were consistent with the notion that at least for some of the commensal strains tested, their high tolerance to SET-M33 is linked to the degradation of the compound in cultures.

### Section 4: Testing the efficacy of SET-M33 in an in vitro infection model

The experimental data described above suggested a complex picture of how SET-M33 could be used to target porcine respiratory pathogens: on the one hand, the high SET-M33 sensitivity of the pathogenic *G. parasuis* strain tested here (compared to several tested commensal nasal microbiota members, **Figure 3**) provides a potential therapeutic opportunity (i.e. targeting of a pathogen while keeping the commensal microbiota intact). On the other hand, the inactivation of SET-M33 by some of these commensals *in vitro* (see **Figure 4**) may limit its efficacy within the context of an *in vivo* infection (where we expect both pathogen and commensals to be present). To address this concern *in vitro*, we established a co-culture infection model consisting of a co-culture of *R. nasimurium* UK1-9 (a SET-M33 tolerant commensal strain able to inactivate the compound) and *G. parasuis* Nagasaki (a SET-M33 sensitive pathogenic strain), initially inoculated in BHI media at a ratio of 1:100 (*Rothia* to *Glaesserella*) to mimic a severe infection (see schematic in **Figure 5**). In the control cultures (no SET-M33 treatment), the co-culture was dominated by *G. parasuis* Nagasaki. In contrast, in SET-M33-treated cultures, *R. nasimurium* UK1-9 was able to grow, while *G. parasuis* Nagasaki was effectively inhibited (**Figure 5B, C**). Thus, these data suggested that SET-M33 retains its selective antimicrobial activity against pathogenic *G. parasuis* even in the presence of commensal microbiota members with the ability to inactivate the compound.

**Figure 5.**
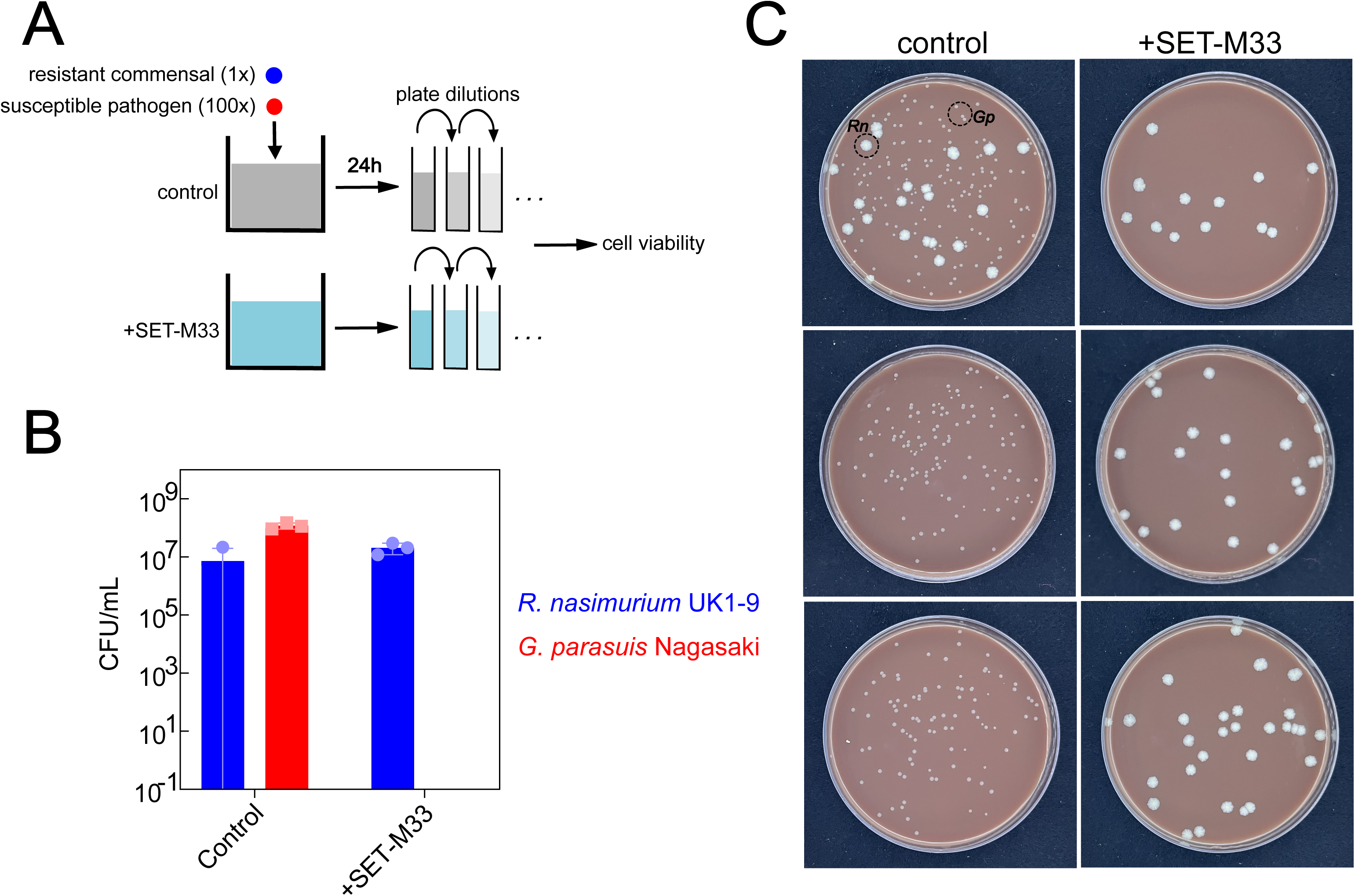
Impact of SET-M33 on pathogen-commensal *in vitro* co-cultures. **A)** Schematic of approach. **B)** Colony forming units (CFUs) of *G. parasuis* Nagasaki (red, pathogenic strain) and *R. nasimurium* UK1-9 (blue, commensal strain) after co-cultivation for 24h in BHI+ either in absence (left) or presence (right) of 10 μM SET-M33. Error bars denote standard deviation (n = 3), circles denote individual replicate cultures. **C)** Corresponding agar plate images used to determine CFUs (shown: 10^-5^ diluted cultures plated on chocolate agar).

## Discussion

As we are facing an era of increasing numbers of bacterial pathogens being resistant to most conventional antibiotics in use today, there is a pressing need to identify suitable use cases for the growing arsenal of newly developed antimicrobial alternatives. In this case study, we focused on one such compound, the synthetic antimicrobial peptide SET-M33, and examined its ability to target three porcine respiratory pathogens, namely *S. suis*, *A. pleuropneumoniae* and *G. parasuis*. Here, we tested the efficacy of SET-M33, and its D-stereoisomer SET-M33D, in *in vitro* cultivation experiments that not only covered pathogenic strains, but also a collection of porcine nasal microbiota members isolated from healthy piglets (34) to examine the potential collateral damage caused by these compounds on the commensal microbiota (38). Our experiments revealed that even within such a comparatively simple *in vitro* model system, there is substantial complexity that emerges from the interplay between SET-M33, pathogens and commensals.

First, we observed large differences in sensitivity to SET-M33 and SET-M33D both among pathogenic (see **Figure 1**) and commensal (see **Figure 2**) strains. These differences in sensitivity did not map neatly to the expectation based on the known differences in bacterial properties (i.e. cell wall composition, see **Figure 3**). For example, although SET-M33D was developed to specifically target gram-positive species such as *S. aureus* (37), it was ineffective against several of the Gram-positive strains tested here (including, notably, a porcine *S. aureus* strain isolated from the nasal cavity). Similarly, the two Gram-negative pathogens *G. parasuis* Nagasaki and *A. pleuropneumoniae* 4074 had completely different sensitivity patterns to both SET-M33 and SET-M33D. These results are in line with previous studies showing that the prediction of target pathogens for antimicrobials remains a challenge (42–44). Future efforts may use the results presented here to guide the development of e.g. SET-M33 variants with differential target ranges.

Second, we found that some of the commensal strains tested here were able to inactivate SET-M33 *in vitro*. Inactivation of antimicrobials by microbiota members is likely not restricted to this specific compound, or antimicrobial peptides in general. For example, previous work has already shown that SET-M33 is more susceptible to bacterial proteases than SET-M33D (37), and degradation of antimicrobial peptides by proteases is likely a general resistance mechanism in many bacterial species (45,46). In fact, chemical drug modification by microbiota members is not restricted to bacteria-targeting compounds. For example, recent work has shown that human gut microbiota members can metabolize many orally administered drugs (47), and it is conceivable that such processes are not restricted to the gut microbiota. Future efforts may examine how widespread the microbial modification of antimicrobials and other drugs is within the respiratory microbiota of animals and humans.

Third, despite this inactivation of SET-M33 by some commensals, we found that this compound was still able to selectively eliminate the tested pathogen (i.e. *G. parasuis* Nagasaki) in co-cultures that contained both the pathogen and the SET-M33-inactivating commensal *R. nasimurium* UK1-9 (**Figure 5**). How can we explain this seemingly paradoxical result? A likely explanation are the different time scales at which killing and SET-M33 inactivation occur: SET-M33 (like other antimicrobial peptides) has been shown to act rapidly, perforating the cell membrane of susceptible cells in minutes (32). Therefore, a likely explanation for our observations is that this compound exerts its effect on susceptible bacteria before other community members can inactivate it. Conversely, these results also suggest that degradation of an antimicrobial by the commensal microbiota does not necessarily preclude its use against pathogens. Future efforts may examine how the different time scales of antimicrobial action and degradation can be exploited to selectively target pathogens while minimizing long-term effects on the microbiota.

This study has several limitations. First, here we focused on a comparatively simple model system consisting of reference pathogenic strains and a previously established collection of commensal nasal microbiota members. Therefore, we cannot exclude the possibility that at least some of the observed differences in sensitivity to SET-M33 are strain-, rather than species-dependent. Future efforts may expand on this work, for example by examining to which extent the sensitivity to SET-M33 in each porcine nasal microbiota species is conserved across isolates (both for pathogens and commensals).

Second, in this study we focused on the role of the commensal nasal microbiota, but did not take the host itself into account. It is conceivable that especially in the case of antimicrobial peptides such as the one tested here, host factors such as extracellular proteases may affect the activity of such compounds. Moreover, the commonly observed toxicity of many antimicrobial peptides in host cells (48,49) may limit their therapeutically useful concentration range *in vivo*. As a valuable intermediate step towards *in vivo* testing of SET-M33 against porcine respiratory pathogens, future efforts may use *in vitro* host models, such as porcine nasal organoids (50), to examine the interplay between host, pathogens, commensals, and SET-M33 experimentally.

Third, in this study we only examined the impact of short-term treatments of porcine pathogens and commensals with SET-M33, and did not examine the potential long-term impact due to the emergence of resistance mutations. Although previous studies have suggested that antimicrobial peptides are less prone to generate acquired resistance (8,26,27), it is clear that such resistance can in principle occur (28,29). Future efforts may use this work as a starting point to examine the emergence of such resistance mutations against antimicrobial peptides in both pathogens and commensals.

In conclusion, this study shows that SET-M33 can be used to selectively target pathogenic *G. parasuis in vitro*, and provides a template for identifying new therapeutic use cases for antimicrobial peptides that minimize collateral damage on the commensal microbiota.

## Methods

### Reagents and strains

Unless stated otherwise, all reagents were purchased from Sigma-Aldrich. SET-M33 was provided by SetLance (Italy). Brain Heart Infusion (BHI) medium (53286, Millipore) was supplemented with 0,08 mg/mL NAD (N0632) and 1% (v/v) heat-inactivated pig serum (P9783) and is denoted here as BHI+.

Reference strains of porcine respiratory pathogens used here were: *Glaesserella parasuis* Nagasaki (serovar 5) originally isolated in Japan from a pig with meningitis (51), *Streptococcus suis* P1/7 (serovar 2, gift from Dr. Marcelo Gottschalk, (52)) and *Actinobacillus pleuropneumoniae* 4074 (gift from Dr. Marcelo Gottschalk, (53)). Commensal porcine nasal microbiota strains *Lactobacillus plantarum* KD9-5, *Staphylococcus aureus* EJ41-2 and *Neisseria shayeganii* GM3-3 were originally described in (34); *Glaesserella parasuis* F9 in (54), *Rothia nasimurium* UK1-9 and *Streptococcus pluranimalium* LG3-6 in (55). *Escherichia coli* NCM3722 was originally described in (56). See **supplementary table 1** for a full list of strains used in this study.

### *In vitro* cultivation experiments

#### Bacterial susceptibility test

Susceptibility tests of target bacteria strains were performed in microdilution assay format in 96-well plates. Briefly, SET-M33 was tested in BHI+ using different final concentrations ranging from 0.62 to 20 µM in 2-fold dilutions. Bacterial suspensions were prepared from fresh overnight chocolate agar plate cultures (PolyVitex chocolate agar, Biomerieux, for all strains except *L. plantarum* KD9-5, which was plated on MRS agar) in PBS to OD600 = 0.3 and used to inoculate 96-well plates (Greiner Cat. No 655 180) to a final concentration of 5 x 10^5^ CFU/mL, in a final volume of 100 µL per well, and 100 µL mineral oil (M3516) was added on top to avoid evaporation and condensation (34). After incubation at 37°C in a 5% CO_2_-enriched atmosphere for 24h, OD600 measurements were performed using an automated plate reader (Tecan Nano M+), and absence/presence of growth was confirmed by visual inspection of each plate.

#### Testing residual SET-M33 activity in spent media obtained from commensal strains

Residual SET-M33 activity in media obtained from SET-M33 tolerant commensal strains was determined as follows. Spent media were obtained from BHI+ cultures that were supplemented with 10 µM of SET-M33 (media without SET-M33 addition were used as controls) and inoculated 1:50 with an OD600 = 0.3 suspension from each individual commensal strain (i.e. *R. nasimurium* UK1-9, *S. aureus* EJ41-2 and *S. pluranimalium* LG3-6). Cultures were incubated overnight at 37°C with orbital shaking at 250 rpm. After incubation, cells were removed from cultures by centrifuging for 20 minutes at 4000 rpm and filtering with sterile filters (Whatman Puradisc 30, 0.2 µm pore size). To restore depleted nutrients, fresh BHI+ was added 1:1 (yielding a maximal SET-M33 concentration of 5 µM). These spent media were then used to grow the SET-M33 sensitive strain *Escherichia coli* NCM3722 (56) in 96-well plates (inoculating each culture 1:50 using PBS suspensions at OD600 = 0.3 and a final volume of 150 µL) and 100 µL mineral oil was again added on top to avoid evaporation and condensation. Plates were incubated overnight at 37°C with 5% CO_2_, and bacterial growth was quantified by OD600 measurements (Tecan Nano M+ plate reader).

#### In vitro co-culture model

Bacterial suspensions of *R. nasimurium* UK1-9 and *G. parasuis* Nagasaki were prepared in PBS to a final OD600 = 0.3 as mentioned above. In a 96-well plate, bacterial suspensions were used to inoculate BHI+ with or without 10 µM of SET-M33 in a final volume of 150 µL. Strains were co-inoculated, using a ratio of 1:100 for *R. nasimurium* UK1-9 and *G. parasuis* Nagasaki, respectively, and plates were incubated without shaking at 37°C in with 5% CO_2_ for 24h. After incubation, cultures were serially diluted in PBS, 100 µL of each dilution were plated on chocolate agar, and plates were incubated at 37°C 5% CO_2_ for 48h. The number of counted colonies of each strain was used to assess colony forming units (CFU).

#### Data processing

Unless stated otherwise, experimental data were processed in Excel (Microsoft), with custom MATLAB scripts (Mathworks, version R2021a), and GraphPad Prism (GraphPad Software Inc, version 8.3.0).

## Supporting information

Supplementary information is provided as a single document containing:

– **Supplementary table 1**: list of strains used in the study
– **Supplementary figure 1**: Day-to-day reproducibility of MIC determination
– **Supplementary figure 2**: Control experiment showing that SET-M33 does not get inactivated by bacteria-free BHI medium.

## Acknowledgements

We thank Marcelo Gottschalk for kindly providing the bacterial strains *Streptococcus suis* P1/7 and *Actinobacillus pleuropneumoniae* 4074.

## Author contributions

Conceived and designed the study: AL, MCC, KK. Performed experiments and analyses: AL, OR, KK. Supervised analyses: CF, AP, VA, MCC, KK. Wrote manuscript with contributions from all authors: AL, KK.

