## Supplementary Text for "SET-M33 peptide as a selective *in vitro* antimicrobial agent against the porcine respiratory pathogen *Glaesserella parasuis*"

### Supplementary tables

#### Supplementary table 1. List of strains used in study.

| **Strain** | **Type** | **Source** |
| --- | --- | --- |
| *Streptococcus suis* P1/7 | Pathogenic | (1) |
| *Actinobacillus pleuropneumoniae* 4074 | Pathogenic | (2) |
| *Glaesserella parasuis* Nagasaki | Pathogenic | (3) |
| *Lactobacillus plantarum* KD9-5 | Commensal | (4) |
| *Streptococcus pluranimalium* LG3-6 | Commensal | (5) |
| *Rothia nasimurium* UK1-9 | Commensal | (5) |
| *Glaesserella parasuis* F9 | Commensal | (6) |
| *Neisseria shayeganii* GM3-3 | Commensal | (4) |
| *Staphylococcus aureus* EJ41-2 | Commensal | (4) |
| *Escherichia coli* NCM3722 | Sentinel strain | (7) |

### Supplementary figures

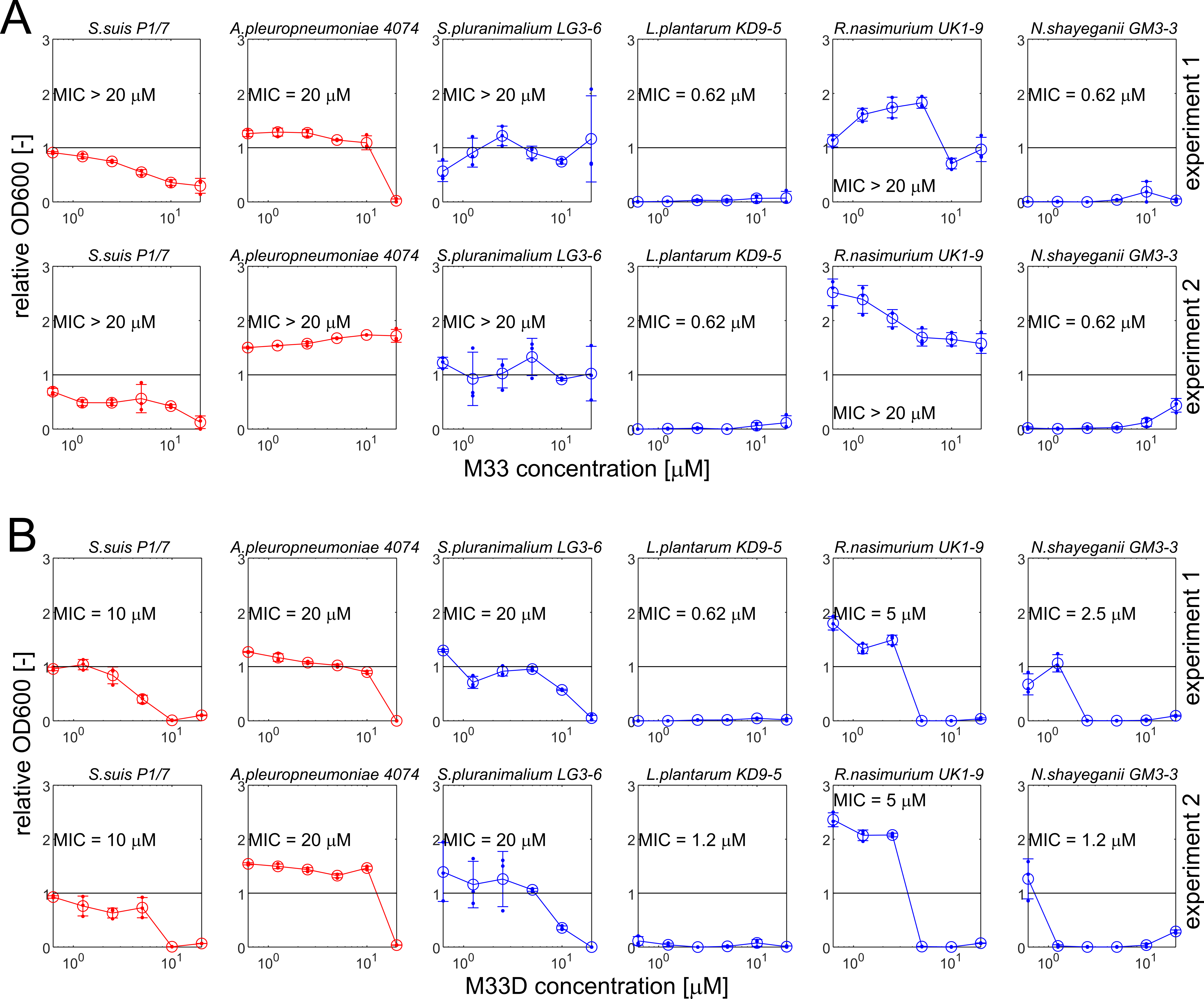

**Supplementary figure 1**. **Day-to-day reproducibility of MIC determination for SET-M33 (A) and SET-M33D (B).** Data shown are from two independent experiments performed on different days (top and bottom row in each panel). Relative OD600 (relative to untreated controls) after 24h incubation in BHI+ with different compound concentrations for each strain. Error bars denote standard deviation (n = 3), small circles denote individual replicate wells. In each case, the MIC was determined as the lowest compound concentration with a relative OD600 of <10%, and the absence of growth was confirmed by visual inspection of the culture plates. Red: selected reference pathogenic strains. Blue: selected commensal strains.

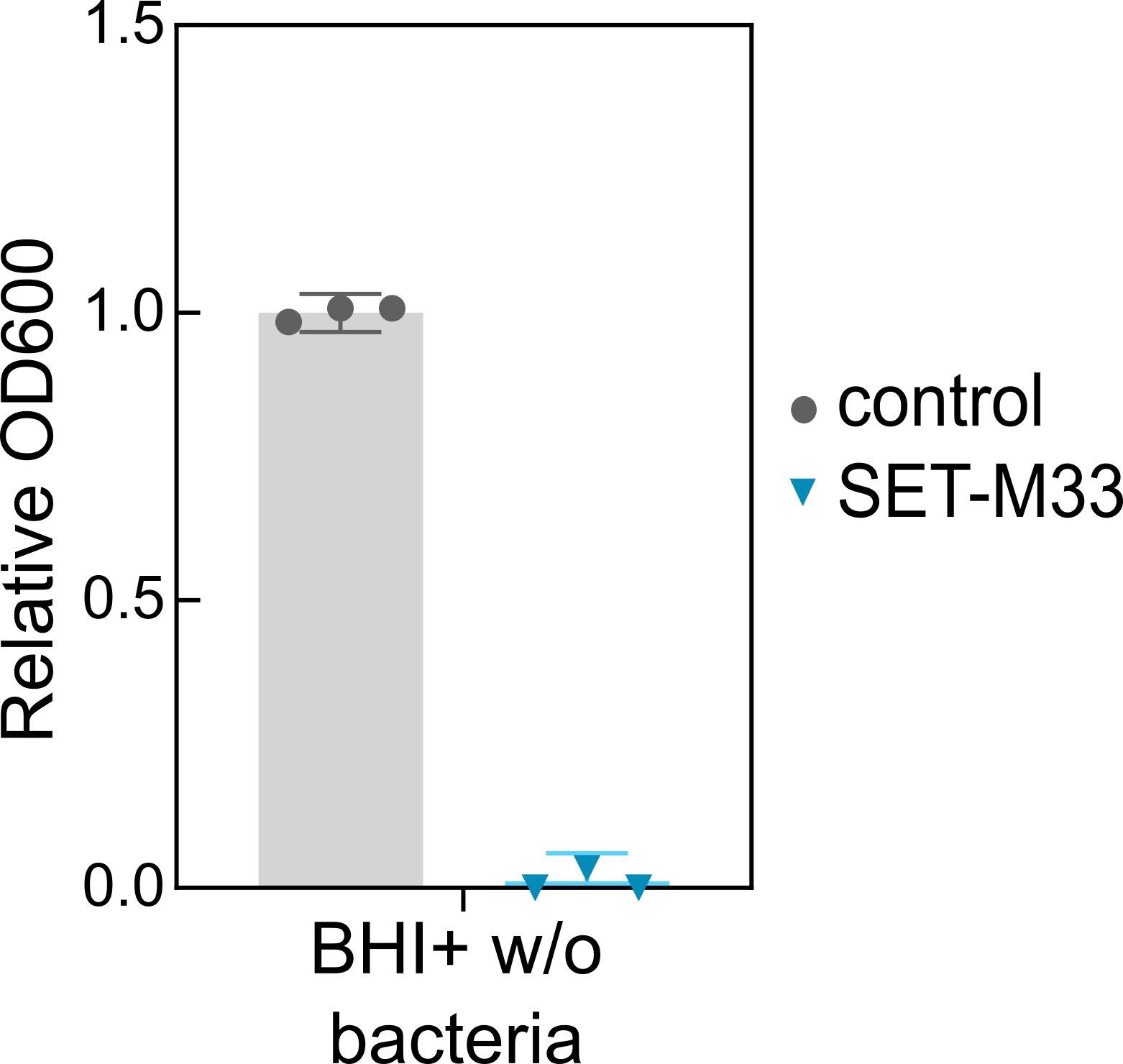

**Supplementary figure 2.** **SET-M33 does not get inactivated by prolonged (i.e. 24h) incubation in BHI+ in absence of bacteria.** Relative OD600 (relative to untreated controls) of SET-M33 susceptible sentinel strain (*E. coli* NCM3722) in cultivation media (BHI+) incubated either without (control) or with 10 μM SET-M33 at 37ºC for 24h. Error bars denote standard deviation (n = 3), small circles denote individual replicate wells.
